# Dopamine D1 and D4 receptors contribute to light adaptation in ON-sustained retinal ganglion cells

**DOI:** 10.1101/2020.10.29.361147

**Authors:** Michael D. Flood, Erika D. Eggers

**Author notes:** Corresponding author: Erika Eggers, Departments of Physiology and Biomedical Engineering, P.O. Box 245051, University of Arizona, Tucson, AZ, 85724, United States of America. Email Addresses: Michael Flood.

## Abstract

The adaptation of ganglion cells to increasing light levels is a crucial property of the retina. The retina must respond to light intensities that vary by 10-12 orders of magnitude, but the dynamic range of ganglion cell responses covers only ∼3 orders of magnitude. Dopamine is a crucial neuromodulator for light adaptation and activates receptors in the D1 and D2 families. D1Rs are expressed on horizontal cells and some bipolar, amacrine and ganglion cells. In the D2 family D2Rs are expressed on dopaminergic amacrine cells and D4Rs are primarily expressed on photoreceptors. However, the roles of activating these receptors to modulate the synaptic properties of the inputs to ganglion cells are not yet clear. Here we used single cell retinal patch-clamp recordings from the mouse retina to determine how activating D1Rs and D4Rs changed the light-evoked and spontaneous excitatory inputs to ON-sustained (ON-s) ganglion cells. We found that both D1R and D4R activation decrease the light-evoked excitatory inputs to ON-s ganglion cells, but that only the sum of the peak response decrease due to activating the two receptors was similar to the effect of light adaptation to a rod-saturating background. The largest effects on spontaneous excitatory activity of both D1R and D4R agonists was on the frequency of events, suggesting that both D1Rs and D4Rs are acting upstream of the ganglion cells.

**New and Noteworthy:** Dopamine by bright light conditions allows retinal neurons to reduce sensitivity to adapt to bright light conditions. It is not clear how and why dopamine receptors modulate retinal ganglion cell signaling. We found that both D1 and D4 dopamine receptors in photoreceptors and inner retinal neurons contribute significantly to the reduction in sensitivity of ganglion cells with light adaptation. However, light adaptation also requires dopamine independent mechanisms that could reflect inherent sensitivity changes in photoreceptors.

## Introduction

Over the course of a day our eyes are exposed to lighting conditions that vary in intensity by 10-12 orders of magnitude (1). In contrast, retinal ganglion cells can only modulate their firing rates by a factor of ∼100-1000 (2). In order to maintain a proper signaling response to light, the retina must constantly tune its sensitivity to ambient luminance conditions. The multiple processes involved in this retinal tuning are collectively referred to as light adaptation.

Studies on light adaptation have found that typically ganglion cell light sensitivity and receptive field size are reduced by increased background illumination (2-7). However, light adaptation can have variable impacts depending on the ganglion cell type (8, 9) and can reduce synchronization of ganglion cell inputs (10) or even change response polarity (11-14) depending on luminance intensity. Because the retina consists of at least 40 classes of ganglion cells (15-18), each with a particular set of conductances and connectivity, it is possible that light adaptation imparts changes to retinal encoding that are unique to each of these individual populations.

For more than half a century visual neuroscientists have known that dopamine is integral to retinal light adaptation, but the full scope of its sites of action and neuromodulatory effects remain inadequately understood and quantified (19, 20). In the retina, dopamine is produced by a specific population of dopaminergic amacrine cells that are located within the inner nuclear layer (19, 21). When the retina is exposed to increased light levels, dopamine is released by dopaminergic amacrine cells into the retina (22, 23). Dopamine mediates its effects primarily in a paracrine manner by binding to dopamine type-D1, -D2, and -D4 receptors located on retinal neuronal subtypes (24-29). Dopamine type D1 receptors (D1Rs) are expressed on horizontal cells and select populations of bipolar, amacrine and ganglion cells (24, 25). Dopamine type D4 receptors (D4Rs) are primarily located on photoreceptors (both rods and cones), but there is also evidence for their expression in some inner nuclear and ganglion cell populations (26, 28, 30, 31). Dopamine type D2 receptors are mainly expressed by dopaminergic amacrine cells, where they act as autoreceptors to limit dopamine release (28, 29, 32). Although the effects of dopamine and its agonists on ganglion cells have been partially characterized (30, 33-39), most studies to date have focused on spike rate modulation in eyecup preps or in dissociated ganglion cells. However, this approach cannot differentiate the complex circuit dynamics that occur upstream of ganglion cells.

In this study, we isolated excitatory currents to ON sustained (ON-s) ganglion cells, which reflect glutamate release by ON bipolar cells. By targeting this synapse, the differential effects of D1R and D4R signaling pathways on light adaptation-induced changes to ganglion cell excitatory inputs could be determined. In addition, the site(s) of action presynaptic or postsynaptic to ganglion cells could be identified. Thus, the results presented here extend our understanding of dopaminergic modulation of retinal function.

## Methods

### Retinal slice and whole mount preparation

Animal protocols conformed with the ARVO Statement for the Use of Animals in Ophthalmic and Visual Research and were approved by the University of Arizona Institutional Animal Care and Use Committee. As previously described (40), C57BL/6J male mice (Jackson Laboratories, Bar Harbor, ME) 35–60 days of age were euthanized using carbon dioxide. The eyes were enucleated, the cornea and lens removed, and the eyecup was incubated in cold extracellular solution (see Solutions and drugs) with 800 U/ml of hyaluronidase for 20 min. The eyecup was washed with cold extracellular solution, and the retina was removed. For slice preparation, the retina was trimmed into an approximate rectangle and mounted onto 0.45-µm nitrocellulose filter paper (Millipore, Billerica, MA). The filter paper containing the retina was then transferred to a hand chopper, sliced into 250-µm-thick slices, rotated 90°, and mounted onto glass coverslips using vacuum grease. Retinal whole mounts were prepared according to the methods outlined by Ivanova et al. (41). Briefly, a hydrophilized cell culture filter insert (Millipore Sigma, Burlington MA) was trimmed down with a Dremel tool to a height of ∼1mm. The retina was cut into 4 equal quadrants and mounted onto our trimmed cell culture insert as needed, one quadrant per ganglion cell experiment. Quadrants were mounted photoreceptor side down by applying slight negative pressure with a custom-trimmed 1 mL syringe. All dissections and light response recording procedures were performed under infrared illumination to preserve the light sensitivity of our preparations.

### Solutions and drugs

Extracellular solution used as a control bath for dissection and whole-cell recordings was bubbled with a mixture of 95% O2-5% CO2, which set the pH to ∼7.4 and contained the following (in mM): 125.00 NaCl, 2.50 KCl, 1.00 MgCl2, 1.25 NaH2PO4, 20.00 glucose, 26.00 NaHCO3, and 2.00 CaCl2. The intracellular solution in the recording pipette used for monitoring excitatory ganglion cell currents contained the following (in mM): 120.00 CsOH, 120.00 gluconic acid, 1.00 MgCl2, 10.00 HEPES, 10.00 EGTA, 10.00 tetraethylammonium-Cl, 10.00 phosphocreatine-Na2, 4.00 Mg-ATP, 0.50 Na-GTP, and 0.1% sulforhodamine-B dissolved in water and was adjusted to pH 7.2 with CsOH. With these concentrations, the driving force for Cl− was calculated as 60 mV in all solutions.

The partial D1R agonist SKF-38393 (SKF, 20 µM; Tocris, Bristol, United Kingdom) was used to selectively activate D1Rs, and the D4R agonist PD-168077-maleate (PD, 500 nM; Sigma) was used to selectively activate D4Rs. Each was diluted in extracellular solution to the given concentration and applied to the bath during the recordings by a gravity-driven superfusion system (Cell Microcontrols, Norfolk, VA) at a rate of ∼1 ml/min. For D4R agonist experiments, as DMSO was initially used to solubilize the drug, our perfusate had a final working DMSO concentration of less than 0.0025% by mass. Chemicals were purchased from Sigma-Aldrich (St. Louis, MO), unless otherwise indicated.

### Whole-cell recordings

All light response recordings were first performed on retinal slices or whole-mounts in a dark-adapted state, followed by recordings performed under drug-added or light-adapted conditions (see Light Stimuli). Retinal slices on glass coverslips or whole-mount preps were placed in a custom chamber and heated to 32° by a TC-324 temperature controller coupled to an SH-27B inline heater (Warner, Hamden CT). For D1R/D4R agonist experiments, dark-adapted light responses were measured first, followed by a 5-min incubation period with the agonist, after which light responses were again measured in the presence of continuous agonist perfusion. For light adaptation experiments, dark-adapted light response measurements were followed by 5 min of light adaptation, before again recording light responses (with background illumination maintained.) Whole-cell voltage-clamp recordings were made from ON-s ganglion cells in retinal slices or whole-mount preparations. Light-evoked (L-) and spontaneous (s) excitatory post-synaptic currents (EPSCs) were recorded from ON ganglion cells voltage clamped at -60 mV, the reversal potential calculated for Cl^-^ currents. For all recordings, series resistance was uncompensated. Electrodes were pulled from borosilicate glass (World Precision Instruments, Sarasota, FL) using a P97 Flaming/Brown puller (Sutter Instruments, Novato, CA) and had resistances of 3–7 MΩ. Liquid junction potentials of 20 mV, calculated with Clampex software (Molecular Devices, Sunnyvale, CA), were corrected before recording. Light responses and spontaneous events were sampled at 10 kHz and filtered at 6 kHz with the four-pole Bessel filter on a Multi-Clamp 700B patch-clamp amplifier (Molecular Devices) before being digitized with a Digidata 1140 data acquisition system (Molecular Devices) and Clampex software.

ON-s ganglion cells were targeted by their large soma size (>15 µM diameter) (42) and identified by their ON-sustained excitatory response to a 500ms-duration light stimulus. Confirmation of ganglion cell morphology and presence of an axon was done at the end of each recording using an Intensilight fluorescence lamp to visualize sulforhodamine-B fluorescence and images were captured with a Digitalsight camera controlled by Elements software (Nikon Instruments, Tokyo, Japan).

### Light stimuli

Full-field light stimuli were evoked with a light-emitting diode (LED; HLMP-3950, λpeak = 525 nm; Agilent, Palo Alto, CA) that was calibrated with an S471 optometer (Gamma Scientific, San Diego, CA) and projected through the camera port of the microscope and onto the stage via a 4x objective. The stimulus intensities were chosen to cover both rod-dominant (9.5, 95.0, 950.0 photons·µm^−2^·s^−1^) and cone-dominant (9.5·10^3^, 9.5·10^4^, and 9.5·10^5^ photons·µm^−2^·s^−1^) response ranges. These intensities were calculated to be equivalent to 4.75, 47.50, 475.00, 4.75·10^3^, 4.75·10^4^, and 4.75·10^5^ R*·rod^−1^·s^−1^, respectively (43). In some locations light intensity is described in R*, which uses the duration of the light stimulus (30 ms). Sequential light responses were recorded with a stimulating interval of 30 s. Stimulus intensities, background rod-saturating light (950 photons·µm^−2^·s^−1^), and duration (30 ms) were controlled with Clampex software by varying the current through the LED. The background intensity was chosen, as it was shown to maximally activate rods (44). A rod-saturating background light was applied for 5 min to light adapt the retina slice or whole-mount prep and was sustained throughout all light-adapted recordings.

### Data analysis and statistics

Between two and four traces of L-EPSCs for each condition were averaged using Clampfit (Molecular Devices). The peak amplitude, charge transfer (Q), time to peak, and decay to 37% of the peak (D37) were determined. The bounds for integration used to calculate Q were marked by the times at which the longest-duration response began and when it returned to baseline, typically 1–2 s. These integration bounds were kept constant for all light responses recorded from the same cell. The time to peak was calculated as the temporal difference between stimulus onset and the response peak amplitude. Because the decay time could not be easily fitted with either a single or double exponential curve, the D37 was determined by computing the time it took for the EPSC to decline from its peak amplitude to 37% of its peak amplitude.

For L-EPSCs, data from each cell were normalized to the maximum response recorded; typically this maximum occurred under dark-adapted conditions at the highest intensity (9.5·10^5^ photons·µm^−2^·s^−1^). If there was no discernable response for a given light intensity after filtering and averaging, the peak amplitude was recorded as 0, and it was excluded from our analysis of response kinetics. Comparisons between experimental conditions and luminance intensities were made with two-way repeated-measures (RM) ANOVA tests using the Student-Newman-Keuls (SNK) method for pairwise comparisons in SigmaPlot (Systat Software, San Jose, CA). If any data were shown to have a non-normal distribution or unequal variance, tests were repeated on the log10 values (or square root values for peak amplitudes) of data. L-EPSC data from retinal slice and retinal whole-mount preparations were compared. Although the absolute values of responses were larger for the data from the whole-mount preparations, when normalized to the maximum value for each cell there was no significant difference in the relationships (slice vs. whole-mount: Q dark-adapted P=0.49, Q after D1R agonist P=0.15, Q after D4R agonist P=0.74; 2-way repeated measures ANOVA). Therefore, slice and whole-mount data were combined.

For spontaneous currents, events were included in the analysis if they occurred after a light response had returned to baseline, but before the 1 s baseline preceding the subsequent light stimulus. Frequency, amplitude, interevent interval (IEI), and decay constants were calculated using MiniAnalysis software (Synaptosoft, Fort Lee, NJ). Decay constants were fit with a single exponent. Effects of treatments on sEPSCs were analyzed at the single cell level with Kolmogorov-Smirnov (K-S) tests in Clampfit. Amplitude, decay τ, and IEI histogram distributions were normalized to the number of events. Average effects of a D1R agonist, a D4R agonist, or light adaptation on sEPSCs in the same cells were analyzed with paired t-tests, and comparisons between different groups of cells were analyzed with unpaired t-tests, after normalizing each cell to its dark-adapted state. Individual cells were only included in the analysis if they had 10 or more spontaneous events per treatment condition. For all studies, differences were considered significant when P ≤ 0.05. All data are reported as means ± 95% confidence interval.

## Results

### Light adaptation reduces L-EPSCs and sEPSCs in ON-s ganglion cells

To assess the contributions of D1R- and D4R-dependent signaling to light adaptation we first had to quantify the effects of a fixed background luminance on ON-s ganglion cell EPSCs. For our light adaptation protocol we chose a background intensity equal to 950 photons·µm^-2^·s that saturates the rod response (44). The light adaptation protocol (Fig. 1A) significantly reduced both the peak amplitude (Fig. 1B) and Q (Fig. 1C) of L-EPSCs in ON-s ganglion cells (2-way ANOVA, peak amplitude p<0.001; Q p<0.001) at all three intensities tested (SNK, p<0.001). Additionally, light adaptation significantly increased time to peak (2-way ANOVA, p=0.001, Fig. 1D) but decreased the decay to 37% of the peak (D37, 2-way ANOVA, p<0.001, Fig. 1E) at all intensities tested (SNK, p<0.001).

**Figure 1.**
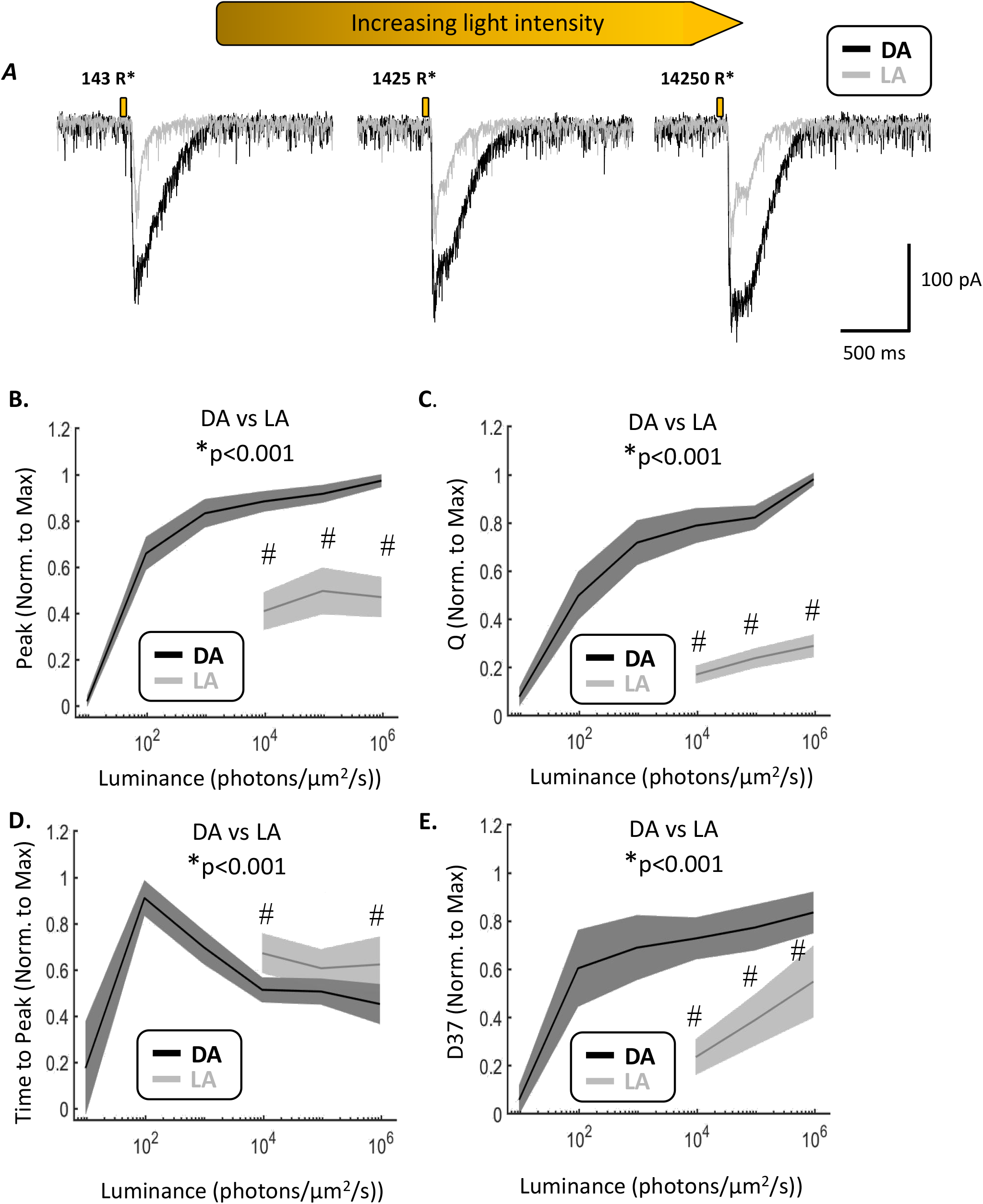
Light adaptation decreases the magnitude and kinetics of L-EPSCs in ON-s ganglion cells. **A**. Example L-EPSC traces from the same cell in response to stimuli of three different light intensities before (black, dark-adapted, DA) and after (gray, light-adapted, LA) light adaptation. 30-ms light stimuli are represented by small gold bars. Stimulus intensities in total rod photoisomerizations are given above each set of traces. **B-E**. Average peak amplitude (B), Q (C), time to peak (D) and D37 (E) of normalized L-EPSCs before (black) and after (gray) light adaptation. 2-way ANOVA main effects of adaptation p values are reported at the top of each graph. * = significant main effects difference between dark and light-adapted states. # = significant pairwise difference between dark and light-adapted states at individual light intensities. Shaded regions represent 95% confidence intervals. For all of these experiments, n = 17 cells from 12 animals.

To determine whether these declines in ON-s ganglion cell L-EPSCs are attributable to pre- or post-synaptic effects, concurrent changes in sEPSC frequency or amplitude were measured (Fig. 2). Analysis of cumulative histograms showed that light adaptation caused significant shifts in sEPSCs towards smaller amplitudes (Fig. 2B) and longer inter-event intervals (Fig. 2C). Light adaptation caused a large decline in average sEPSC frequency in each cell (−61.4% ± 7.4%, paired t-test p<0.00001) as well as a slight decline in amplitude (−19.5 ± 4.3%, paired t-test p=0.00044). This suggests that light adaptation modulates the activity of circuits upstream of ganglion cells, and may induce post-synaptic changes to ganglion cell receptors. Because we did not pharmacologically isolate true miniature EPSCs for these experiments, it should be noted that the decline in sEPSC amplitude might also be attributable to changes in upstream activity.

**Figure 2.**
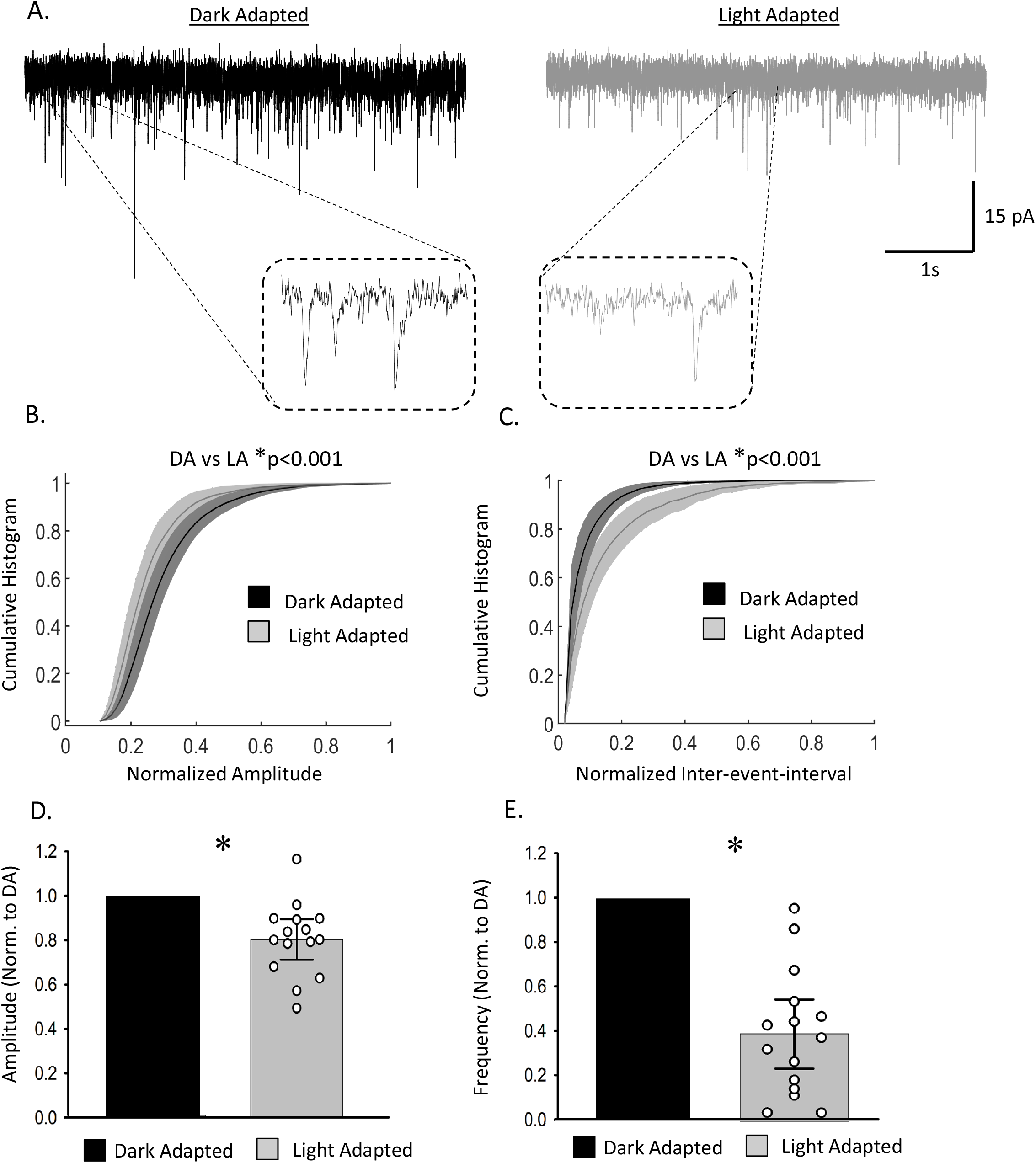
Light adaptation decreases the frequency and amplitude of sEPSCs in ON-s ganglion cells. **A**. Example sEPSC traces from the same cell before (black) and after (gray) light adaptation. **B-C**. Average cumulative histogram of sEPSC amplitudes (B) and inter-event intervals (C) before and after light adaptation. Shaded regions represent 95% confidence intervals. Bin frequencies were normalized to total number of events recorded under each condition, and amplitude/IEI values were normalized to maximum values recorded on a cell-by-cell basis. 2-way ANOVA main effects p-values for DA vs LA are reported above each graph. **D-E**. Average changes in sEPSC amplitude (D) and frequency (E) after light adaptation. Values were normalized on a cell-by-cell basis to that recorded under dark-adapted conditions. Individual data points are marked by empty white circles. Error bars = +/-standard error. For all of these experiments, n = 15 cells from 10 animals.

### D1R activation reduces the magnitude of L-EPSCs in ON-s ganglion cells

Given that light adaptation significantly modulates ON-s ganglion cell L-EPSCs, the contributions of D1Rs and D4Rs were then determined. First, the effect of a D1R agonist on ON-s L-EPSCs was measured (Fig. 3A). L-EPSC peak amplitudes (Fig. 3B) and Q (Fig. 3C) were significantly diminished (2-way ANOVA, peak amplitude p<0.001; Q p=0.009) with significant pairwise differences at the top 4 intensities for peak amplitudes (SNK, p<0.05) and at 9.5·10^4^ photons·µm^-2^·s^-1^ for Q (SNK, p=0.04), although some variability was noted (shown by 95% confidence intervals). D1R agonist application significantly increased time to peak values (2-way ANOVA, p=0.015, Fig. 3D) with a significant pairwise difference at 950 photons·um^-2^·s^-1^ (SNK, p=0.013), but did not have a significant impact on D37 values (2-way ANOVA, p=0.079, Fig. 3E). However, light adaptation caused significantly greater reductions to peak amplitude and Q values (2-way ANOVA, peak p=0.002, Q p<0.001) than D1R activation (Fig. 3B, C). Light-adapted L-EPSCs also had significantly longer time to peak values and significantly shorter D37 values (2-way ANOVA, T2P p<0.001, D37 p=0.03, Fig. 3D, E) than D1R L-EPSCs. These results suggest that while D1R application has a significant impact on L-EPSCs, it does not fully recapitulate the effects of light adaptation.

**Figure 3.**
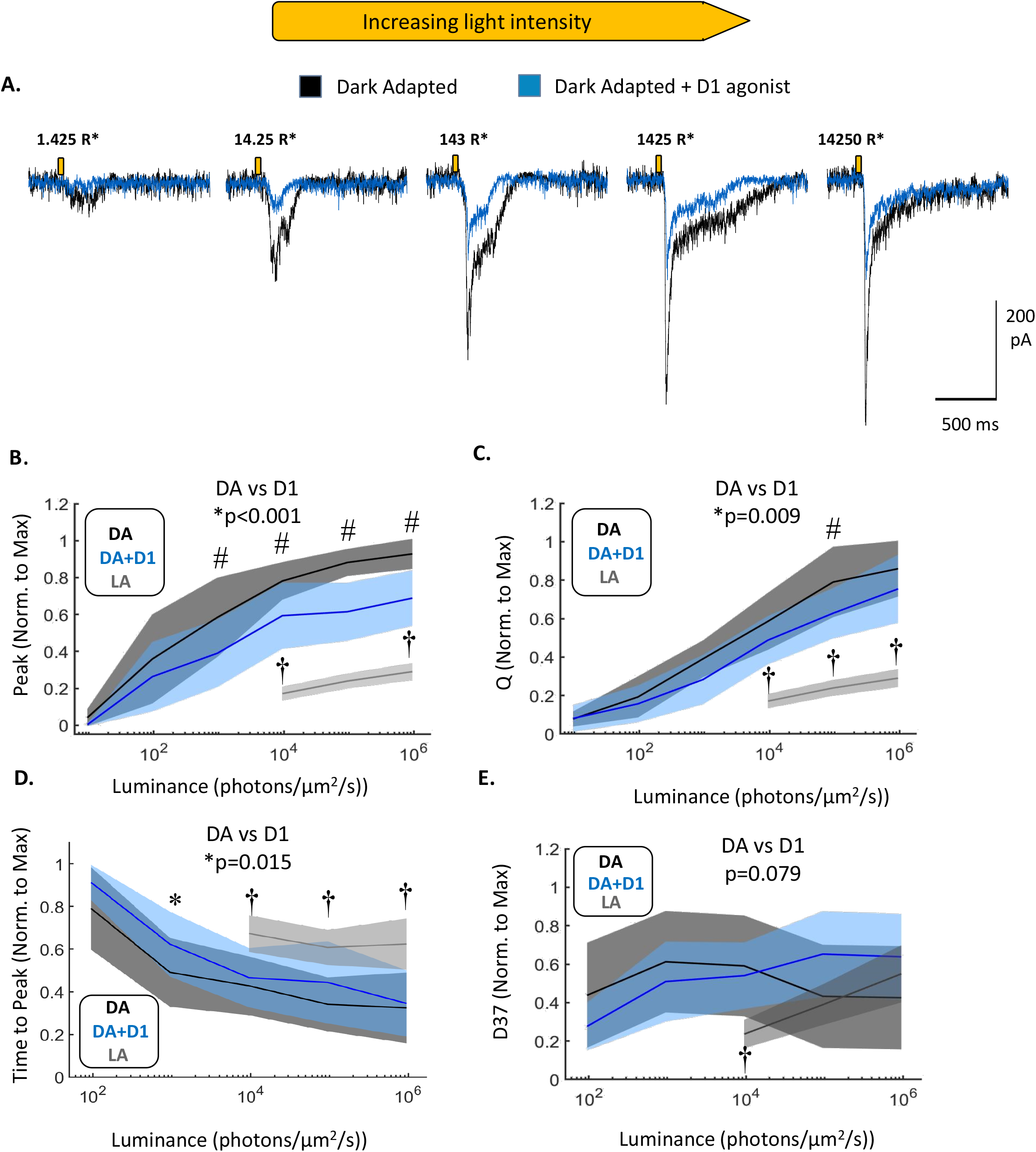
D1R activation decreases the magnitude and times to peak of L-EPSCs in ON-s ganglion cells. **A**. Example L-EPSC traces from the same cell at increasing light intensities before (black) and after (blue) application of the D1R agonist SKF-38393 (20 µM). The gold bars represent 30 ms light stimuli. Stimulus intensities in total rod photoisomerizations are given above each set of traces. **B-E**. Comparison of average peak amplitude (B), Q (C), time to peak (D) and D37 (E) between dark-adapted (DA) and D1R-agonized (DA+D1) conditions. All values are normalized on a cell-by-cell basis to the maximum values of each parameter recorded for each cell. Average normalized values of light-adapted ganglion cells from figure 1 are included for reference. Main effects of agonist treatment p-values for two-way ANOVAs are shown above each graph. * = significantly different main effect between dark-adapted and D1R agonist-treated state. # = significantly different pairwise comparison between dark-adapted and D1R agonized states. † = significantly different pairwise comparison between D1R agonized and light-adapted states. Shaded regions represent 95% confidence intervals. For all of our D1R L-EPSC experiments, n = 9 cells from 8 animals.

### D1R activation reduces frequency and amplitude of sEPSCs in some but not all ON-s ganglion cells

We next examined the effects of D1R activation on sEPSCs in the cells that light responses were recorded from (Fig. 4A). Analysis of cumulative histograms showed that D1R activation significantly shifted sEPSC amplitudes (Fig. 4B) and inter-event intervals (Fig. 4C) towards smaller and longer values, respectively. Although the sEPSC distributions were shifted significantly, average sEPSC amplitude and frequency values after D1R treatment for each cell were smaller, (Fig. 4D, E), but not to a significant degree (paired t-test, amplitude p=0.46; frequency p=0.125, n = 5 cells from 5 animals). The magnitude of D1R-mediated effects on average sEPSC amplitude and frequency varied, and one cell showed increased amplitude and frequency after D1R application. When we compared amplitude and inter-event interval distributions on a single cell level, we found that D1R activation caused significant shifts towards smaller amplitudes and longer inter-event intervals in 3/5 cells (K-S, p<0.05). We found no correlation between the degree of changes in sEPSC amplitudes or frequencies and changes in L-EPSCs. When we compared the normalized changes induced by D1R activation and light adaptation to average sEPSC amplitude and frequency, we found no significant differences. Owing to variability in this data, it is unclear whether light adaptation mediates its effects on sEPSCs via D1Rs.

**Figure 4.**
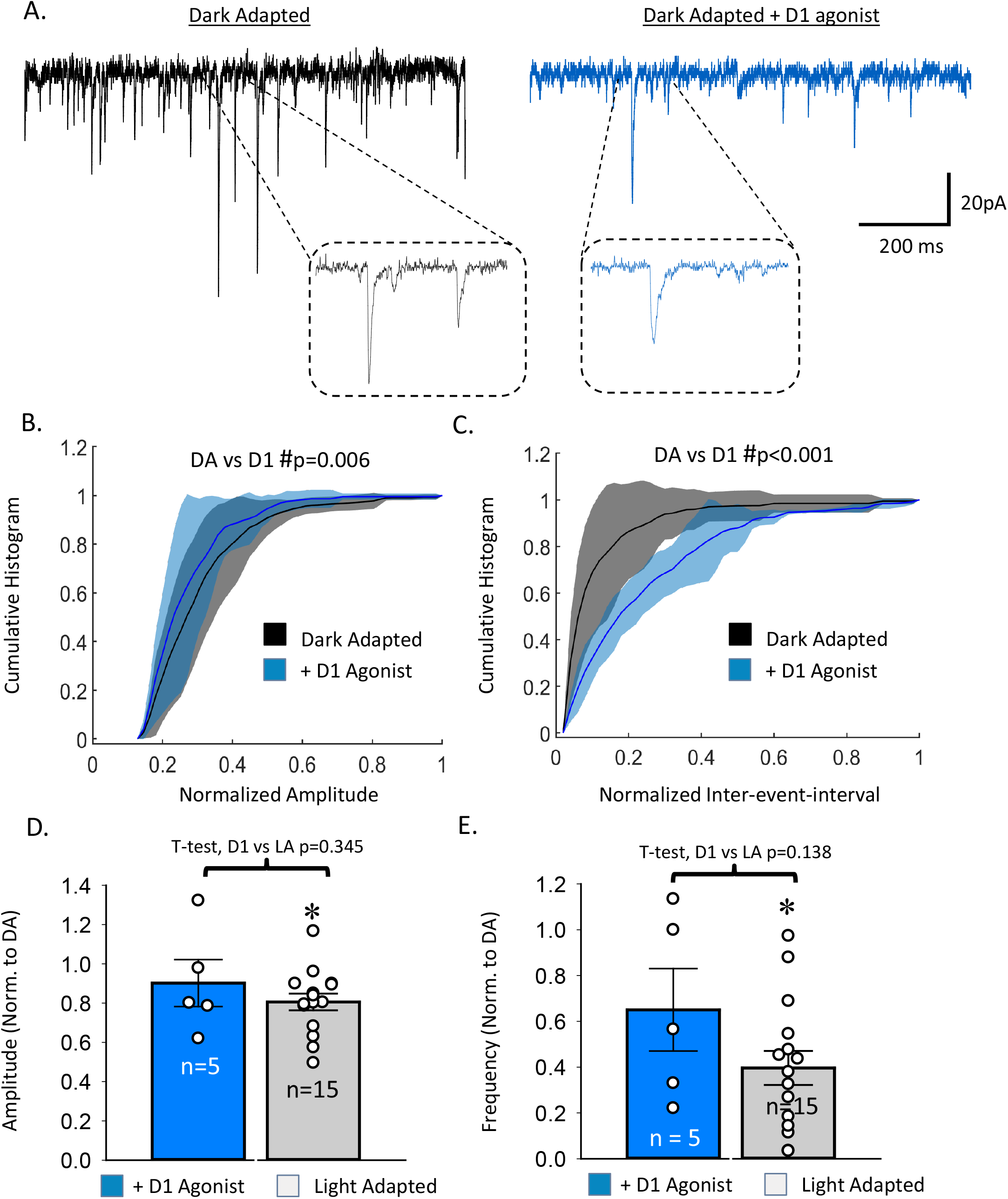
D1R activation decreases the frequency of sEPSCs in some but not all ON-s ganglion cells. **A**. Example sEPSC traces from the same cell before (black) and after (blue) application of the D1R agonist SKF 38393 (20 µM). **B-C**. Average cumulative histogram of sEPSC amplitudes (B) and inter-event intervals (C) before and after D1R activation. Shaded regions represent 95% confidence intervals. Bin frequencies were normalized to total number of events recorded under each condition, and amplitude/IEI values were normalized to maximum values recorded on a cell-by-cell basis. 2-way ANOVA main effects p-values for DA vs D1R are reported above each graph. **D-E**. Average changes in sEPSC amplitude (D) and frequency (E) after D1R agonist application or light adaptation (included from Figure 2 for reference). Average values for each treatment were normalized on a cell-by-cell basis to that recorded under dark-adapted conditions. Individual data points are marked by empty white circles. Error bars = +/-standard error. # = significant difference between dark-adapted and D1R agonist conditions. * = significant difference between D1R agonist and light-adapted conditions. For all D1R sEPSC experiments, n = 5 cells from 5 animals.

### D4R activation decreases L-EPSCs in ON-s ganglion cells

Similar to our analysis of the effects of a D1R agonist, we examined the effects of a D4R agonist on light-evoked currents in ON-s ganglion cells (Fig. 5A). Both peak amplitude (Fig. 5B) and Q (Fig. 5C) were significantly decreased by D4R activation (2-way ANOVA, p<0.001, n=7 cells from 7 animals), with significant pairwise differences at 950, 9.5·10^4^ and 9.5·10^5^ photons·µm^-2^·s^-1^ for peak amplitudes and at 9.5·10^3^, 9.5·10^4^, 9.5·10^5^ photons·µm^-2^·s^-1^ photons for Q (SNK, p<0.05) and some variability. No significant difference in time to peak (Fig. 5D) or D37 (Fig. 5E) between dark-adapted and D4-agonist treated responses (2-way ANOVA, time to peak p=0.904; D37 p=0.719, n=6 cells from 6 animals) were identified. However, similar to the data on D1R activation, light adaptation caused significantly greater reductions to both peak amplitude and Q than D4R activation (2-way ANOVA, peak amplitude p<0.001, Q p<0.001, Fig. 5B, C). Light-adapted L-EPSCs also had significantly longer times to peak and shorter D37 values (2-way ANOVA, T2P p<0.001, D37 p=0.026, Fig. 5D, E) than L-EPSCs after activating D4R. This data suggests that while D4R application has significant impacts on L-EPSCs, it cannot fully account for changes that occur during light adaptation.

**Figure 5.**
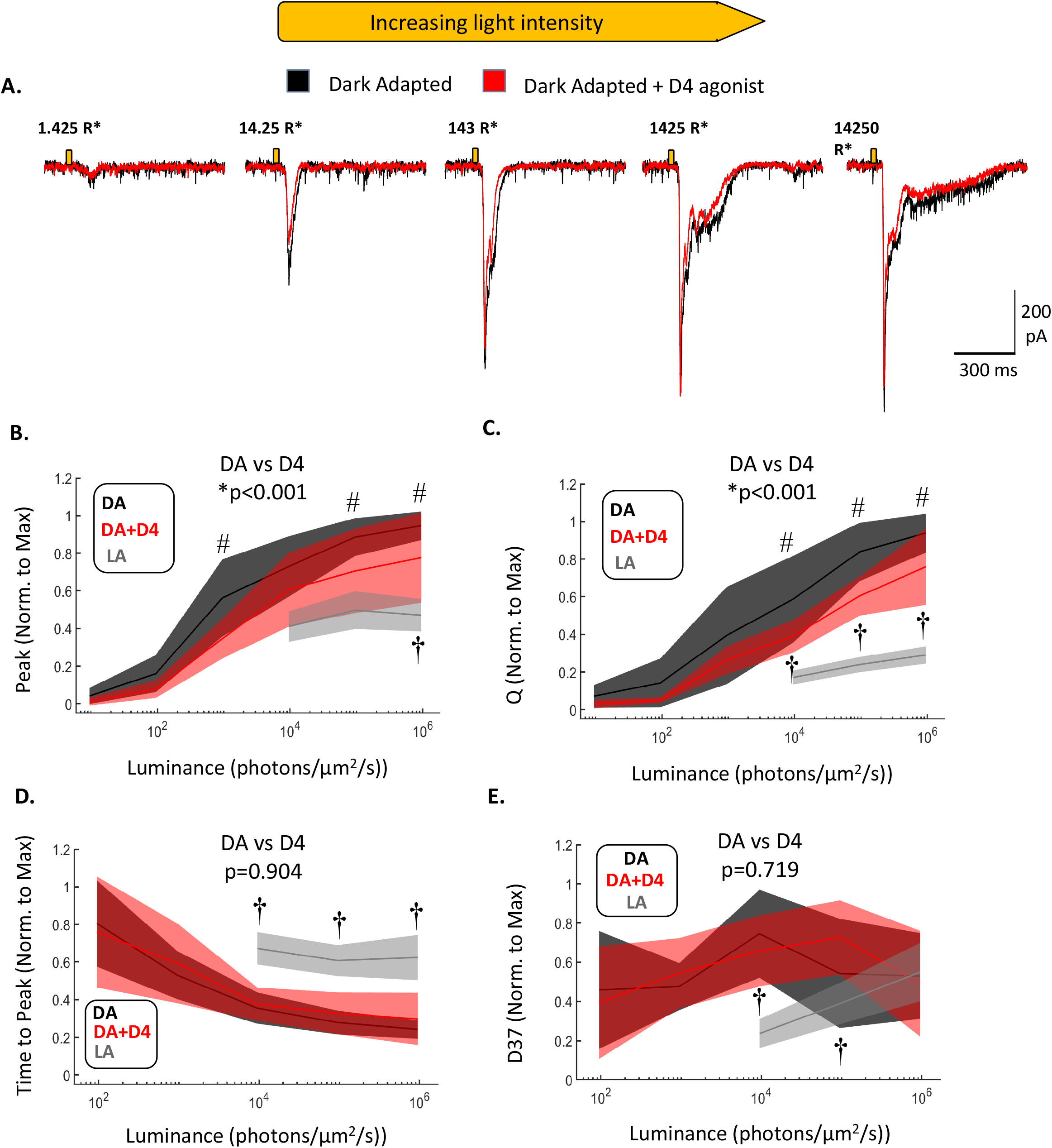
D4R activation decreases the magnitude but not kinetics of L-EPSCs in ON-s ganglion cells. **A**. Example L-EPSC traces from the same cell at increasing light intensities before (black) and after (red) application of the D4R agonist PD-168077 maleate (500 nm). The gold bars represent 30 ms light stimuli. Stimulus intensities in total rod photoisomerizations are given above each set of traces. **B-E**. Comparison of Q (B) peak amplitude (C), time to peak (D) and D37 (E) between dark-adapted (DA) and D4R-agonized (DA+D4) conditions. Light-adapted (LA) values from figure 1 are included for comparison. Main effects of agonist treatment p-values for two-way ANOVAs are shown above each graph. * = significantly different main effect between dark-adapted and D4R agonist-treated states at each light intensity. # = significantly different pairwise comparison between dark-adapted and D4R agonist states. † = significantly different pairwise comparison between D4R agonist and light-adapted states. Shaded regions represent 95% confidence intervals. For the D4R L-EPSC experiments, n = 7 cells from 6 animals.

### D4R activation decreases the frequency and amplitude of sEPSCs in ON-s ganglion cells

We next analyzed the impact of D4R activation upon ON-s ganglion cell sEPSCs (Fig. 6A). Cumulative histograms showed that D4R activation significantly shifted sEPSCs towards smaller amplitudes (Fig. 6B) and longer inter-event intervals (Fig. 6C). Treatment with a D4R agonist caused significant reductions in the average amplitude (Fig. 6D) and frequency (Fig. 6E) of sEPSCs in ON-s ganglion cells (paired t-test, frequency p=0.00369; amplitude p=0.00353). On the single cell level, D4R activation caused significant shifts in amplitudes towards smaller values, and inter-event intervals towards longer values, in 4/5 cells (K-S, p<0.01). This suggests pre-synaptic changes in tonic bipolar cell release and potential post-synaptic modifications to receptors on ganglion cells. Again, we found no correlation between the magnitude of D4R activation’s impact on sEPSC and L-EPSC parameters. There were no significant differences between the normalized changes induced by D4R activation and light adaptation to average sEPSC amplitudes and frequencies. Collectively, this data suggests that light adaptation mediates its effects on sEPSCs via D4Rs.

**Figure 6.**
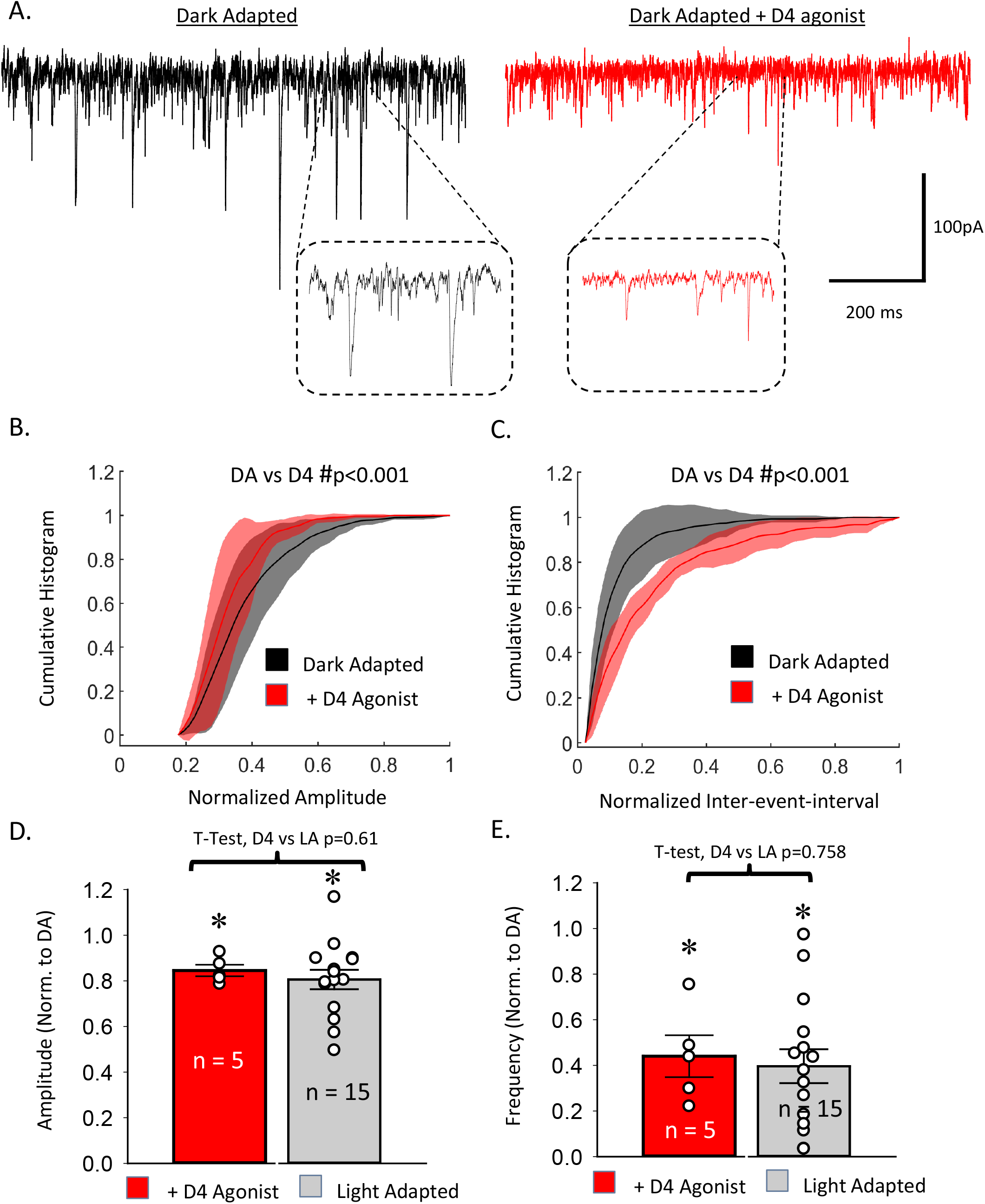
D4R activation decreases the frequency of sEPSCs and slightly diminishes sEPSC amplitudes in ONs ganglion cells. **A**. Example sEPSC traces from the same cell before (black) and after (red) application of the D4R agonist PD-168077 maleate (500 nM). **B-C**. Average cumulative histogram of sEPSC amplitudes (B) and inter-event intervals (C) before and after D4R activation. Shaded regions represent 95% confidence intervals. Bin frequencies were normalized to total number of events recorded under each condition, and amplitude/IEI values were normalized to maximum values recorded on a cell-by-cell basis. 2-way ANOVA main effects p-values for DA vs D4R agonist are reported above each graph. **D-E**. Average changes in sEPSC amplitude (D) and frequency (E) after D4R agonist application or light adaptation. Average values for each treatment were normalized on a cell-by-cell basis to that recorded under dark-adapted conditions. Individual data points are marked by empty white circles. Error bars = +/-standard error. # = significant difference between dark-adapted and D4R agonist conditions. * = significant difference between D4R agonist and light-adapted conditions. For these experiments, n = 5 cells from 4 animals.

### Combined effects of D1R/D4R activation approximate some but not all changes to L-EPSCs and sEPSCs that are induced by light adaptation

To see whether the effects of light adaptation on ON-s ganglion cell light responses could be fully explained by dopamine receptor activation we created a simple additive model for our L-EPSC peak amplitude and Q values (Fig. 7A, B). For peak amplitude values (Fig. 7A), the hypothetical combination of D1R/D4R agonist treatment (dashed gold line) closely approximated the measured response reduction caused by light adaptation (grey line). However, this was not the case for our Q data (Fig. 7B) where our additive model did not cause nearly the same reduction as light adaptation. We also created an additive model for our sEPSC amplitude and frequency values (Fig. 7C, D) showing the hypothetical effects of D1R/D4R agonist co-activation. In this case, the model produced similar declines to average sEPSC peak amplitudes (Fig. 7C), but decreased average sEPSC frequency to a greater degree than light adaptation (Fig. 7D). These models suggest that D1R and D4R activation may have synergistic effects on these parameters, or perhaps that there are additional changes induced by light adaptation that act through dopamine-independent mechanisms.

**Figure 7.**
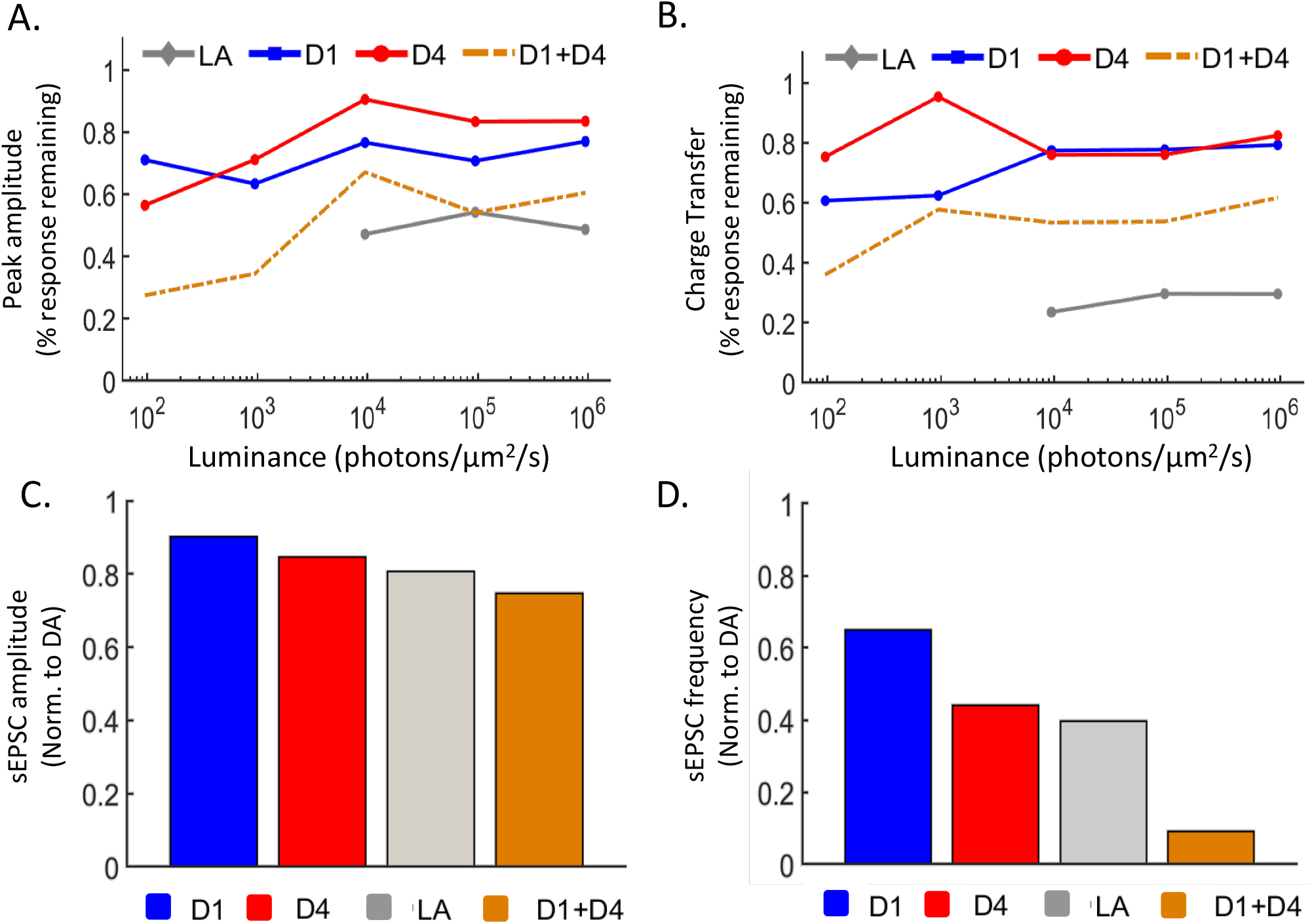
Additive effects of dopaminergic signaling do not fully explain light adaptation. **A-B**. Average percent of response remaining (as compared to dark-adapted conditions) after light adaptation (LA, gray), application of a D1R agonist (D1, blue), or D4R agonist (D4, red), as well as the theoretical reduction one would witness from the additive effects of D1R-D4R agonist co-application (D1+D4, dashed line). The additive effects of D1R/D4R agonist co-application approximate the effects of light adaptation in diminishing peak amplitude (A), but do not match the decline in charge transfer witnessed upon light adaptation (B). **C-D**. Average % of sEPSC amplitude (C) and frequency (D) compared to dark-adapted conditions after light adaptation (LA, grey), D1R activation (D1, blue), and D4R activation (D4, red). The hypothetical impact of combined D1R/D4R activation is also shown (D1+D4, gold). The combined effects of dopamine agonists approximate the effect of light adaptation on average sEPSC amplitude (C), but not average sEPSC frequency (D).

## Discussion

In this study, we examined the effects of light adaptation and dopamine receptor agonists on excitatory inputs to ON-s ganglion cells, in order to gauge the relative contribution of dopaminergic pathways to luminance adaptation. Light adaptation reduced the magnitude of L-EPSCs while delaying their times to peak and speeding up their decays. This correlated with a decrease in the frequency and amplitude of sEPSCs. Activation of either D1Rs or D4Rs replicated a portion of these changes but did not recapitulate them fully. The measured effects of light adaptation on excitatory inputs to ON-s ganglion cells largely fit with prior data. Previous reports have demonstrated reduced light sensitivity (3-6, 45), as well as faster response kinetics (13, 46) of ganglion cell spike rates in the light-adapted retina. Previous reports also showed that light adaptation significantly reduces resting ganglion cell spike rates (8, 47), which could be mediated by the reductions in spontaneous currents measured here. Because dopamine is released during light adaptation and modulates retinal activity via activation of both D1Rs and D4Rs (19, 20), it makes sense that treatment with either a D1R or D4R agonist only partially induces the changes seen upon light adaptation.

### Activation of D1Rs can modulate retinal output at multiple synaptic sites

Activating D1Rs decreased L-EPSCs magnitude and slightly increased time to peak (Fig. 3). Our analysis of D1R activation’s effects on sEPSC inter-event intervals suggested that most of these changes are attributable to decreased ON bipolar cell output rather than direct effects upon ON-s ganglion cells. Upstream of ON-s ganglion cells there are multiple sites where D1R activation should modify retinal activity. In the outer plexiform layer, photoreceptor activity should not be directly modified, as they do not express D1Rs (24, 25, 48). However, activation of D1Rs in mouse horizontal cells reduces voltage-gated calcium currents (49), which should reduce negative feedback onto photoreceptors (50, 51) and result in enhanced glutamate release (52). This should decrease depolarization of ON bipolar and ganglion cells, in agreement with our results here. Along with their effects on photoreceptor release, there is evidence that horizontal cells can potentially act via feed-forward GABA_A_ synapses to modify ON cone bipolar cell output (53-58). A recent study in rabbits found evidence for a D1R-mediated increase in GABA_A_ receptor expression in the dendrites of ON cone bipolar cells (59) that led to increased surround strength. In the present study, full-field light stimuli were used that should maximally activate the surrounds of upstream ON cone bipolar cells. Thus, if D1R activation increases the strength of ON cone bipolar cell surrounds, application of a D1R agonist should result in decreased ON cone bipolar cell responses to light, which would also help to explain our result of reduced excitatory input to ON-s ganglion cells.

It is also likely that some populations of ON cone bipolar cells are directly affected by D1R activation. Farshi et al. (24) demonstrated D1R expression in ON type 5-2, 6 and 7 bipolar cells, all of which likely form synapses onto ON-s ganglion cells (60, 61) with ON type 6 bipolar cells forming the majority (62). Previous research has demonstrated a role for D1Rs in reducing voltage-gated currents in ON cone bipolar cells (63-65). D1Rs depolarize ON cone bipolar cell resting membrane potentials (59) and modulate GABAergic conductance (66-68), all of which could have significant impacts on excitatory output to ON ganglion cells. While ON cone bipolar cells have not been tested, we have also previously shown that D1R activation can reduce evoked inhibitory inputs to rod bipolar cells (69) and OFF cone bipolar cells (70, 71), possibly by inhibiting amacrine cell calcium currents as in horizontal cells (49). A potential reduction of inhibition to ON cone bipolar cells would not be expected to decrease their outputs onto ON-s ganglion cells. However, a general reduction of inhibition could strongly affect inputs to ganglion cells due to inhibitory connections between amacrine cells (40, 72) such as the recently suggested amacrine cell population that receives primarily ON cone bipolar cell excitatory input and inhibits AII amacrine cells (73, 74).

### Activation of dopamine type-4 receptors likely reduces glutamate release by photoreceptors

We found that D4R activation significantly reduced L-EPSC magnitude (Fig. 5), with a concurrent decrease in sEPSC frequency (Fig. 6C, E). These changes suggest D4R-induced modulation of both phasic and tonic release of glutamate by ON cone bipolar cells. We also found that D4R activation reduced sEPSC amplitude (Fig. 6D), suggesting a post-synaptic change in ionotropic glutamate receptors that could help explain our findings. However, it is possible that this difference in average sEPSC amplitude could arise from reductions in coordinated vesicle release by ON cone bipolar cells. Overall, our results show that D4R activation reduces excitatory inputs to ON-s ganglion cells. Our results support a previous study in light-adapted rabbit retina that saw increased spontaneous spiking activity in ON-s ganglion cells after inhibiting dopamine type-2 family receptors, suggesting that a D4R agonist would decrease spontaneous spiking (33). However, another study on light-adapted rat retina found that antagonizing D4Rs decreases light sensitivity and maximum firing rate in ON ganglion cells, suggesting that D4R activation should increase both of these parameters (75). Since this study was in light-adapted retina (when D1R- and light-induced changes should be active), instead of the dark adapted retina used in the present study and used ganglion cell spiking as their metric that cannot differentiate between pre- and post-synaptic effects, it is difficult to compare the findings between the studies. There is also evidence for D4R expression in certain ganglion cell populations (30, 32, 35, 36, 76) that could affect ganglion cell excitability (31) that is not measured in the voltage-clamp experiments described here. The predominant location for D4Rs in the retina is on photoreceptors (26, 27, 29, 77-79). Activation of D4Rs diminishes photoreceptor light sensitivity (80) and modifies calcium currents (81-83), with the overall effect of decreasing glutamate release (84). Decreased tonic glutamate release from photoreceptors would likely lead to a sustained depolarization of ON bipolar cells and a diminished capacity for both spontaneous and light-evoked glutamate release onto ON ganglion cells via activity-dependent adaptation mechanisms (85, 86). Together with our data, this suggests that the main location of the D4R effects on ON-s ganglion cells comes from the modulation of photoreceptors.

### The full effects of light adapting mechanisms likely require light

We found that neither D1R nor D4R agonists were able to completely recapitulate the effects of our light adaptation protocol on excitatory ganglion cell currents. Our simple additive model for the proposed effects of D1R/D4R co-activation also failed to do so (Fig. 7), although the true effects of such a treatment may be synergistic rather than additive. However, it is likely that light adaptation also involves dopamine-independent mechanisms. Several studies have identified activity-dependent effects on vesicle supply (87-89), calcium influx (85, 86), glutamate receptor cycling (90, 91) and release probability (10) that modify the behavior of retinal circuits. In addition, light-dependent release of nitric oxide (92, 93) has been implicated as another source of retinal adaptation that can modify the activity of multiple retinal populations (94-96). In our study, we utilized brief light stimuli that were separated by significant periods of darkness to reduce the impact of these mechanisms in the dark-adapted state, but many of these changes were probably induced to some degree by our light-adapting background.

## Conclusion

Light adaptation, although a straightforward concept from the perspective of signal processing, is a phenomenon of intricate complexity when considering the diverse molecular, cellular and circuit mechanisms that are involved. In this study, we assessed the contributions of specific dopaminergic signaling pathways to light adaptation of a relatively well-characterized retinal circuit. We found that activation of either dopamine pathway contributed to light-induced changes via modulation of retinal activity upstream of ON-s ganglion cells, and perhaps via modulation of post-synaptic receptors on ON-s ganglion cells. Although the likely main sites of dopaminergic action of these two pathways are located at different retinal populations, their effects on full-field light-evoked excitatory currents at the ganglion cell level may be similarly brought about through modulation of photoreceptor release dynamics. Future work will need to be done to characterize the differential contribution of dopaminergic signaling to the modulation of retinal interneuron receptive fields and to isolate the actions of these pathways on inner vs. outer retinal activity.

## Acknowledgements

The authors would like to thank members of the Eggers laboratory for helpful comments on this manuscript.

## Funding

This work was supported by the National Institutes of Health [grant numbers RO1-EY026027, 4T32HL007249-40], the National Science Foundation [NSF CAREER award #1552184] U.S. Department of Veterans Affairs (VA RX002615) and the International Retinal Research Foundation.

## Notes

### Competing Interest Statement

The authors have declared no competing interest.

